# How basic components of information-updating interact to encourage variation in the results of empirical studies of within and transgenerational plasticity

**DOI:** 10.1101/2021.04.12.439424

**Authors:** Judy A Stamps, Alison M Bell

## Abstract

Experiences of parents and/or offspring are often assumed to affect the development of trait values in offspring because they provide information about the external environment, but it is currently unclear how information from different sources and times might combine to affect the information-states that provide the foundation for the patterns observed in empirical studies of developmental plasticity in response to environmental cues. We analyze Bayesian models designed to mimic fully-factorial experimental studies of within and transgenerational plasticity (TWP), in which parents, offspring, neither or both are exposed to cues from predators, to determine how different durations of cue exposure for parents and offspring, the devaluation of information from parents or the degradation of information from parents would affect offspring estimates of environmental states related to risk of predation at the end of such experiments. We show that the effects of different cue durations, the devaluation of information from parents, and the degradation of information from parents on offspring estimates are all expected to vary as a function of interactions with two other key parameters of information-based models of TWP: parental priors and the relative cue reliability in the different treatments. Our results suggest empiricists should expect to observe considerable variation across experimental studies of TWP based on simple principles of information-updating, without needing to invoke additional assumptions about costs, tradeoffs, development constraints, the fitness consequences of different trait values, or other factors.

## Introduction

The trait values expressed by the offspring of one generation can be affected by their own experiences earlier in life (within-generational plasticity, WGP), by the experiences of their parents (transgenerational plasticity, TGP), or by the combined effects of both (within and transgenerational plasticity, TWP) [1-5]. It is often assumed that one reason why the experiences of parents and offspring might have adaptive effects on offspring trait values is that those experiences provide information about conditions in the external environment that the offspring are likely to experience later in life [6-12]. In addition, information about conditions in the external environment can also be provided by genes, inherited epigenic factors and parental phenotypes [7, 13, 14]. Hence, in order to appreciate how information from an individual’s distant and immediate ancestors and its own personal experiences might combine to affect the development of traits that are the focus of studies of WGP, TGP or TWP, we must consider how information from a variety of different sources combines within and across generations to affect the information-state of that individual.

In principle, Bayesian updating is the best way to combine information from different sources and different times to estimate the value of variables in the external environment [15]. As a result, in recent years researchers have begun to use Bayesian approaches to study how information from ancestors and personal experiences might combine over the course of ontogeny [16-22], review in [23]. These models assume that an individual’s estimate of conditions in the external environment can change over ontogeny, as its initial estimate based on information from its ancestors (its naïve prior distribution) is updated on the basis of information provided by its own experiences. In turn, changes in an individual’s estimate over time are assumed to drive WGP, because an individual’s estimate of conditions in the external environment (e.g. its estimate of predator density at that locality) is assumed to affect the trait values it develops in response to that estimate (e.g. its level of antipredator behavior). By extension, these models predict that if different individuals with the same naïve prior distribution were exposed to different informative experiences over the course of ontogeny, they would develop different trait values in response to those experiences [16, 17, 20].

By definition, empirical studies of WGP, TGP and TWP focus on phenotypic traits, not information-states. However, when WGP, TGP or TWP occur in response to cues that provide information about the external environment, variation among individuals in their information-states provides the foundation for variation in the trait values observed in experimental studies of these phenomena. As a result, it is important to understand how the basic components of information-based models (e.g. naïve priors, cue reliability, the duration of exposure to different cues, etc.) interact to jointly affect the information-states of individuals. However, it is difficult to do this in information-based models that generate predictions about the trait values of individuals. This is because such models necessarily include assumptions about many factors in addition to an individual’s information-state that can affect the development and expression of its trait values (e.g. costs of sampling, developmental constraints, or the fitness consequences of expressing different trait values)(ibid). As a result, in such models, it is difficult to tell how much information-updating, per se, contributes to their results.

A different way to study how information-updating might affect developmental plasticity is to focus on information-states, instead of phenotypes [19, 21, 22]. Models which consider how information-states change within and vary across individuals as a function of variation in prior distributions, cue reliability and other key components of information-based models have provided insights into a number of topics in developmental biology, including the ways that information-updating contributes to variation across individuals or genotypes in their developmental trajectories [19, 21, 24], and to variation in age-dependent plasticity across individuals and among empirical studies [22]. These analyzes indicate that empiricists can expect to observe a variety of different patterns in empirical studies of these topics, simply based on the ways that we would expect information from different sources to combine over ontogeny within individuals. By extension, these results suggest that it might not be necessary to invoke assumptions about developmental constraints, tradeoffs, costs of sampling, the fitness consequences of trait values, or other factors to account for at least some of variation in results that empiricists have observed in studies of WGP.

Recently Stamps and Bell [25] extended this approach by using simple Bayesian approaches to model experimental studies of TWP in response to informative cues. They found that interactions between two basic components of such models (the parent’s prior at the beginning of the study, and the relative reliability of the cues in the different treatment groups) had profound effects on the patterns of offspring information-states expected at the end of those experiments. Here, we expand these analyses to consider three other variables that are likely to vary across experimental studies of TWP: 1) differences between the duration of exposure to the same cues in parents and offspring, 2) the extent to which information from parents is devalued, as compared to the information based on offspring experience, and 3) the extent to which information from parents is degraded as it is passed from parents to their offspring. As is described below, we consider how variation among experiments in these three variables, in conjunction with variation among experiments in parental priors and the relative reliability of the cues in the different treatments, would affect the patterns of offspring information-states expected at the end of empirical studies of TWP.

As was the case in [25], we illustrate this approach by modelling fully-factorial experiments of TWP in response to cues from predators, in which parents, offspring, both or neither are exposed to cues from predators, and then the trait values of the offspring are measured at the end of the experiment. Where P = exposed to cues from predators and N = not exposed to cues from predators, and where the first letter indicates the parental treatment and the second letter indicates the offspring treatment, the four treatment groups in this type of study consist of NN (neither parents and offspring exposed to cues), PN (parent exposed, offspring not exposed), NP (parent not exposed, offspring exposed) and PP (both parents and offspring exposed).

The results described in [25] were based on assumptions about experimental design and information-updating which are unlikely to be valid in many empirical studies of TWP. First, with respect to experimental design, we assumed that parents and offspring were exposed to the same cues for the same period of time. This assumption is frequently violated in experimental studies of TWP. For instance, of the 13 experimental studies of TWP in response to cues from predators listed in a recent review [26], the duration of exposure to the cues for parents and for their offspring differed in 11 of them. Hence, one important question is whether and how different durations of exposure to the same cues in parents and offspring might contribute to differences among the offspring in TWP experiments in their information states at the end of these experiments.

Second, in [25] we assumed that exposure to the same cues for the same period of time in parents and offspring would provide the same information to offspring. There are at least two reasons why this assumption need not be valid. First, theory indicates that information based on parental experiences should be devalued, relative to information based on offspring experiences, if environmental conditions might change across or within generations [7, 27]. In nature, changes between generations in the value of a state of the environment can occur if offspring develop in a different environment than their parents, e.g. as a result of natal dispersal in spatially heterogeneous environments [11, 14] or as a result of temporal shifts in environmental conditions that may occur between generations [7, 11]. The results of recent studies of experimental evolution in nematodes support the hypothesis that temporal autocorrelation for the environments of parents and offspring encourage the evolution of transgenerational plasticity, by showing that parental experiences had adaptive effects on offspring traits in lineages which evolved when the environmental conditions during the parental and offspring generation were strongly correlated with one another, but not in lineages in which they were not [28, 29].

In addition, if environmental conditions might change within generations, a parent’s estimates of those conditions based on their experiences earlier in life might no longer be accurate by the time that they transmit information about these conditions to their offspring [7, 30]. This possibility is implicitly acknowledged by empiricists who study TWP when they use experimental protocols that minimize the time that elapses between parental exposure to the cues and the time when parents transfer information to their offspring (e.g. when parents are exposed to cues from predators as adults, just before offspring production)[e.g. 31-32].

Second, information from parents might be degraded between the time that parents perceived the cues and the time that parents transferred information based on those cues to their offspring. Degradation of information from parents based on their exposure to a given cue is likely because the transmission of information from parents to offspring requires additional proximate steps that are not required when offspring are directly exposed to the same cue [30]. For instance, parents that detect a cue from predators might produce a signal (e.g. alter their parental behavior) based on their updated estimate of the chances that this predator lives nearby [34]. Then their offspring would need to detect this signal and use it to update their own estimate of the probability that the predator is in residence. Since neither the parent’s production of the signal nor their offsprings’ detection of that signal are likely to be error-free, the information based on the parent’s experience could become less reliable by the time it reaches their offspring. In contrast to the devaluation of information provided by parents, which is assumed to be an adaptive response to spatial and or temporal heterogeneity in environmental conditions, the degradation of the information provided by the parents is assumed to be an unavoidable consequence of the noise introduced into the information pathway by the proximate mechanisms by which parents pass information based on their own experiences along to their offspring.

Hence, in the current study we considered how differences in the duration of exposure to the same cues in parents and offspring and the devaluation or degradation of information from parents would interact with previously-examined variables (parental priors, relative cue reliability) to affect the patterns of offspring estimates expected at the end of the treatments in experimental studies of TWP in response to cues from predators.

## Material and methods

Details about the design and biological rationale for the models used as the basis for those in the current article are provided in previous publications [19, 21, 22, 25]. In brief, we assumed that each parent began with a prior distribution (Prior), based on information from their ancestors (e.g. via genes, inherited epigenetic factors, grand-parental experiences), as well as any informative experiences the parents had before the onset of the experiment. We illustrate the results for three informative parental Prior distributions, with different means (0.1, 0.5 and 0.9), but the same variance (0.04) (For definitions of these and other terms, see Supporting Information).

We assumed that cues from the predator in the P treatment were always informative, and used beta functions with shapes modelled by α > β to indicate the shapes of the cumulative likelihood functions for the conditions in the P treatments. We analyzed two sets of models which differed with respect to their assumptions about the reliability of the information provided by the N treatment. In the first set (N-models), the information provided by the N treatment was less reliable than the information provided by the P treatment. We analyzed this situation by assuming that the information provided by conditions in the P treatment was highly reliable (likelihood with a shape modelled by α = 8, β = 1) while conditions in N provided no information about the state (i.e., the cumulative likelihood function for the N treatment had a uniform distribution (α = 1, β = 1). In the second set of models (N* models), the information provided by the P treatment and the information provided by the N treatment were equally reliable. In this case, the cumulative likelihood function for the N treatment was the mirror-image of the cumulative likelihood function for the P treatment. For instance, if the experiences in the P treatment resulted in a cumulative likelihood function that indicated with a high level of reliability that the value of the state of the environment was likely to be high (e.g. likelihood modelled by a beta distribution with a shape indicated by α = 8, β = 1), conditions in the N treatment had a cumulative likelihood function that indicated with the same level of reliability that the value of the state was likely to be low (e.g. likelihood with a shape modelled by α = 1, β = 8).

For all of the models, we computed the offspring estimates of the value of the state at the end of their respective treatments, where each estimate was the mean of the offspring posterior distribution at the end of the experiment. Hence, in the current article, ‘offspring estimate’ indicates a key component of the offspring’s information-state at the end of the experiment, namely, its best estimate of the value of the state of the environment at that point in time.

### Duration of exposure to the same cue in parents and offspring

Preliminary analyses showed that if parents and offspring were exposed to the same cues for the same period of time, differences between the parental and the offspring generation in the age of onset of the exposure period had no effects on the results. For instance, our analyses indicated that offspring estimates of predator density would be the same if their parents had been exposed for two weeks to cues from predators just before gamete production as if the offspring themselves had been exposed for two weeks to the same cues as juveniles. These results occur because in Bayesian updating models which assume that the true state of the environment is unlikely to change over time, if different subjects with the same prior distribution are exposed to the same cues, the order in which they were exposed to those cues has no effect on their final posterior distributions. Because the order-indifference of Bayesian updating is particularly relevant to analyses of sensitive periods and age-dependent plasticity, we defer discussion of this point to a study of that topic (Stamps, in prep.)

In contrast, preliminary analyses suggested that different durations of exposure to the same cues in parents and offspring might have strong effects on offspring estimates at the end of their respective treatment periods. In order to investigate the effects of different durations of exposure to the conditions P or N for parents and offspring on the results, we divided the total treatment period, T, for each generation into four intervals of equal length, and then specified the likelihood function for exposure to the cue for one interval, and the likelihood function for no exposure to the cue for one interval. For instance, to model a situation in which parents were exposed to the cues for a longer period than the offspring, we assumed that parents in the P treatment were exposed to cues from the predator for all four intervals, whereas the offspring in the P treatment were exposed to the cues from the predator for either one, two or three intervals, and were exposed to no cues from predators for the remaining interval(s). Similarly, to model a situation in which offspring were exposed to the cues for a longer period than the parents, we assumed that offspring in the P treatment were exposed to the cues for all four intervals, but parents in the P treatment were exposed to the cues for one, two or three intervals, and were exposed to no cues for the remaining intervals. In all of these models, the N group was maintained with no cues from predators for all four intervals. We then used Bayesian updating to compute the mean values of the offspring posterior distributions at the end of the offspring treatment period for each of the treatment groups (NN, PN, NP and PP), based on the likelihood functions for the four intervals in the parental generation and the four intervals in the offspring generation.

As in [25], we computed separate models for the N-situation (in which the conditions during one time interval in the P treatment provided much more reliable information than the conditions during one interval in the N treatment) and for the N* situation (conditions during one interval in the P treatment provided information as reliable as the conditions during one interval in the N treatment). For the N-models, we assumed that in a P treatment in which the individual was not exposed to the cues in all four intervals, no information was provided in an interval that lacked the cue. That is, the likelihood function for one interval spent in the absence of the cue had a shape indicated by a uniform distribution (α =1, β = 1). In the N* models, we assumed that in a P treatment in which the individual was not exposed to the cues in all four intervals, the information provided by one interval spent in the absence of the cue was as reliable as the information provided by one interval spent in the presence of that cue. For instance, if the likelihood function for one interval in the presence of cues from a predator had a shape indicated by α = 2.5, β = 1,, the likelihood function for one interval in the absence of those cues had a shape indicated by α =1, β = 2.5.

### Cues based on parental experience are devalued or degraded

Although the devaluation and the degradation of information from parental experiences are assumed to occur as a result of different processes (see Introduction and Discussion), in both situations it is assumed that the information provided by the signal that parents pass along to their offspring based on their exposure to a given cue is less reliable than the information provided by the cue to which the parents had been exposed. In Bayesian terms, this means that although we would expect the likelihood function for the signal provided to their offspring by parents in the P treatment to have the same mean value as the cue in the P treatment for the offspring, we would expect the likelihood function for the parental signal to have a higher variance than the likelihood function for the cues in the P treatment for the offspring. Hence, we used the same procedure to model the devaluation of information from the parents and the degradation of information from the parents. That is, we assumed for both situations that the signal from the parent and the cue for the offspring had likelihood functions with the same mean, but different variances. For instance, instance, if the likelihood function for the P treatment for offspring had a shape modelled by α = 8, β = 1 (mean = 0.89, variance = 0.01), the likelihood function for the signal provided by parents to offspring by the parents in the P treatment might have a shape modelled by α = 3.5, β = 0.44 (mean = 0.89, variance = 0.02). That is, we assumed that exposure to cues from predators in the P treatment yielded the same point estimate of the value of the state of the environment for parents and offspring (in this case, a relatively high value of 0.89, on a scale of 0 to 1), but the reliability of the information provided by the parents to their offspring (indicated by the variance of the likelihood function) was lower than the reliability of the information provided by the same experience for the offspring.

In the N-models, the reliability of the information for the P treatments differed for parents and offspring, as was indicated above. However, given our assumption for these models that the information in the N treatments was unreliable (see above), we assumed that the information provided by the N treatments was equally unreliable for both the parent and the offspring generation (modelled by α = 1, β = 1).

In the N* models, in which conditions in the N treatment provided information as reliable as those in the P treatments, we devalued the information provided by parents in the N treatment groups to their offspring. For instance, if the likelihood function for the P treatment for the offspring had a shape modelled by α = 8, β = 1, the likelihood function for conditions in the N treatment for offspring had a shape modelled by α = 1, β = 8, but the likelihood function for the information provided by parents to their offspring had a shape modelled by α = 0.44, β = 3.5.

All of the other assumptions and parameter values for these models were the same as those for the models described in [25].

## Results

In the ‘baseline’ models, in which parents and offspring were exposed to the same cues for the same period of time, and in which information from parents was neither devalued or degraded, the resulting patterns of offspring estimates varied as a function of the relative reliability of the cues in the different treatments and the parental Priors (Fig 1). In the N-models, in which the information provided by conditions in one treatment (here P, indicating the presence of cues from predators) was much more reliable than the information provided by the conditions in the other treatment (here, N, indicating the absence of cues from predators), we observed a ‘jump-up’ pattern, in which the value for the NN group was low, and the values for the other three groups (NP, PN, PP) were high and similar (but not identical) to each other. However, this pattern only occurred when the state of the environment indicated by the parental Prior differed from the state indicated by the cue in the P treatment. Since we assumed here that the cues in the P treatment indicated that the value of the state of the environment was high, we observed the jump-up pattern when the parental Prior strongly contradicted the cue (Prior mean = 0.1)(Fig 1a). In contrast, when the parental Prior and the cues in the P treatment indicated the same value of the state (Prior mean = 0.9), the offspring estimates were similar for all four treatment groups. This result occurred because in these models, the N treatment had no effect on an individual’s information-state, so the results were primarily affected by the ‘discrepancy rule’ of Bayesian updating (see [24] for more on this topic).

**Fig 1.**
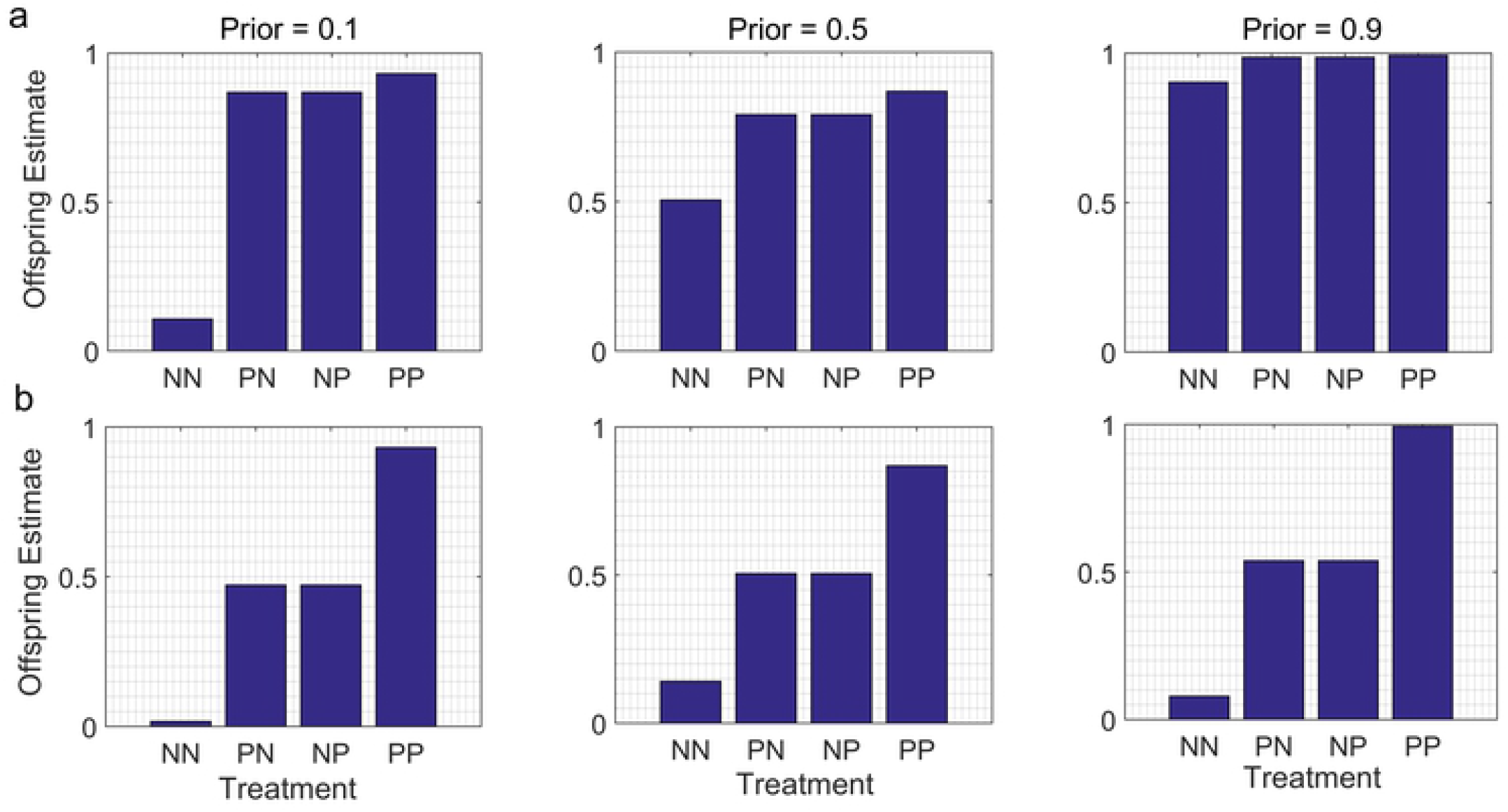
How differences in the reliability of the cues in the different treatments affect offspring estimates of the value of conditions in the external environment. P treatment = exposure to cues from a predator, N treatment = no cues from predator, first letter = parental treatment, second letter = offspring treatment. Predicted offspring estimates at the end of an experiment for each of four treatment groups (NN, PN, NP and PP) are indicated for parental Priors with three different means (0.1, 0.5, 0.9) and the same variance (0.04). a. N-models: conditions in the N treatment provide much less reliable information than conditions in the P treatment. In this example, information provided by conditions in the P treatment is modelled by a cumulative likelihood function with a shape indicated by α = 8, β = 1, indicating that the value of the state is likely to be high; information provided by conditions in the N treatment is modelled by a uniform distribution (α = 1, β = 1). b. N* models. Conditions in the N treatment provide information as reliable as conditions in the P treatment; the likelihood functions for the P and N treatments are mirror-images of one another. In this example, conditions in the P treatment indicate with a high level of reliability that the state of the environment is likely to be high (likelihood modelled by α = 8, β = 1); conditions in the N treatment indicates with equally high reliability that the state of the environment is likely to be low (likelihood modelled by α =1, β = 8).

In the N* models, in which the information provided by conditions in the P treatment and the information provided by conditions in the N treatment were equally reliable, we observed a ‘step-up’ pattern, in which the offspring estimates for the NN treatment were low, those for the PP treatment were high, and those for the NP and PN treatments were intermediate. In this case the pattern did not vary as a function of the parental Prior: for the same set of parameter values, the step-up patterns were similar for a range of parental Priors (Fig 1b). This result occurred because in the N* models, by the end of the experiment reliable information about the state of the environment had been provided by the cues in both treatment groups, so after the offspring were exposed to two doses of informative cues (based upon the parent’s experience and their own personal experience), the initial estimate provided by the parental Prior no longer had much effect on their estimates of the value of the state of the environment. These results have been described and discussed in detail in [25], so we merely present examples of these patterns here for comparison with the results of the models in which we relaxed our assumptions about cue durations, devaluation and degradation.

When parents and offspring were exposed to the same cues for different periods of time, the main effect was to generate differences between the offspring estimates for the PN and NP groups; these differences were not observed in the baseline models (compare Fig 1 with Figs 2 and 3). As intuition would suggest, the offspring estimates were higher for the treatment group which included the generation that had been exposed to the cues for a longer time. For instance, if the offspring in the P treatment group were exposed to the cues for a longer period than their parents, PN < NP (Fig 2a,b). Conversely, if the offspring in the P treatment group were exposed to the cues for a shorter time than their parents, then PN > NP (Fig 3a,b).

**Fig 2.**
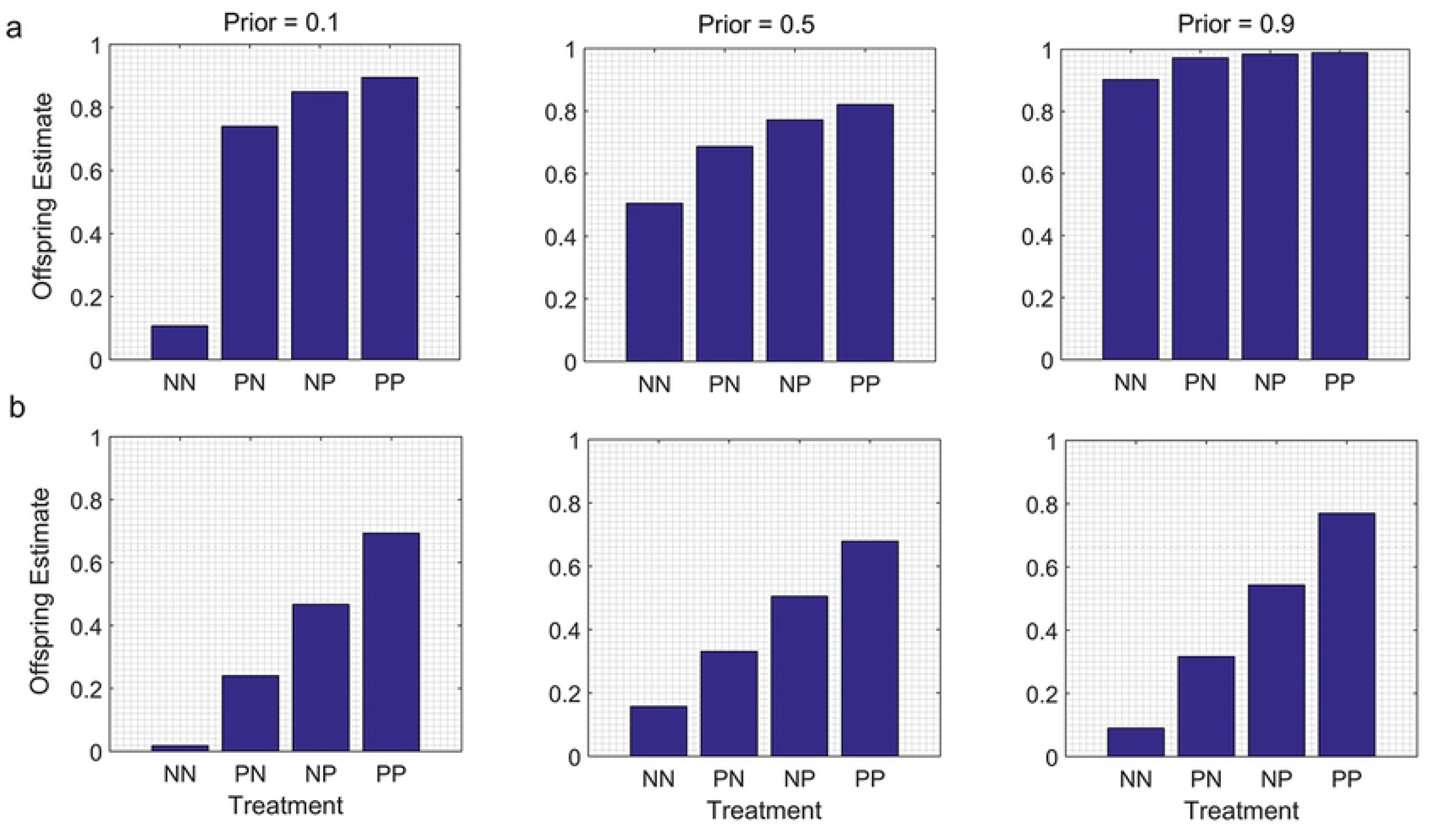
Duration of exposure to the same cues is longer for offspring than for parents. In the P treatment groups, parents are exposed to the presence of cues from predators for two of four time intervals, but offspring are exposed to cues from predators for all four time intervals (see text). a. N-models. In this example, the likelihood function for exposure to cues from a predator for one time interval has a shape indicated by α = 2.5, β = 1; the likelihood function for the absence of cues for one interval has a shape indicated by a uniform distribution (α =1, β = 1). b. N* models. In this example, the likelihood function for exposure to cues from a predator for one interval has a shape indicated by α = 2.5, β = 1; the likelihood function for the absence of cues for one interval has a shape indicated by α =1, β = 2.5.

**Fig 3.**
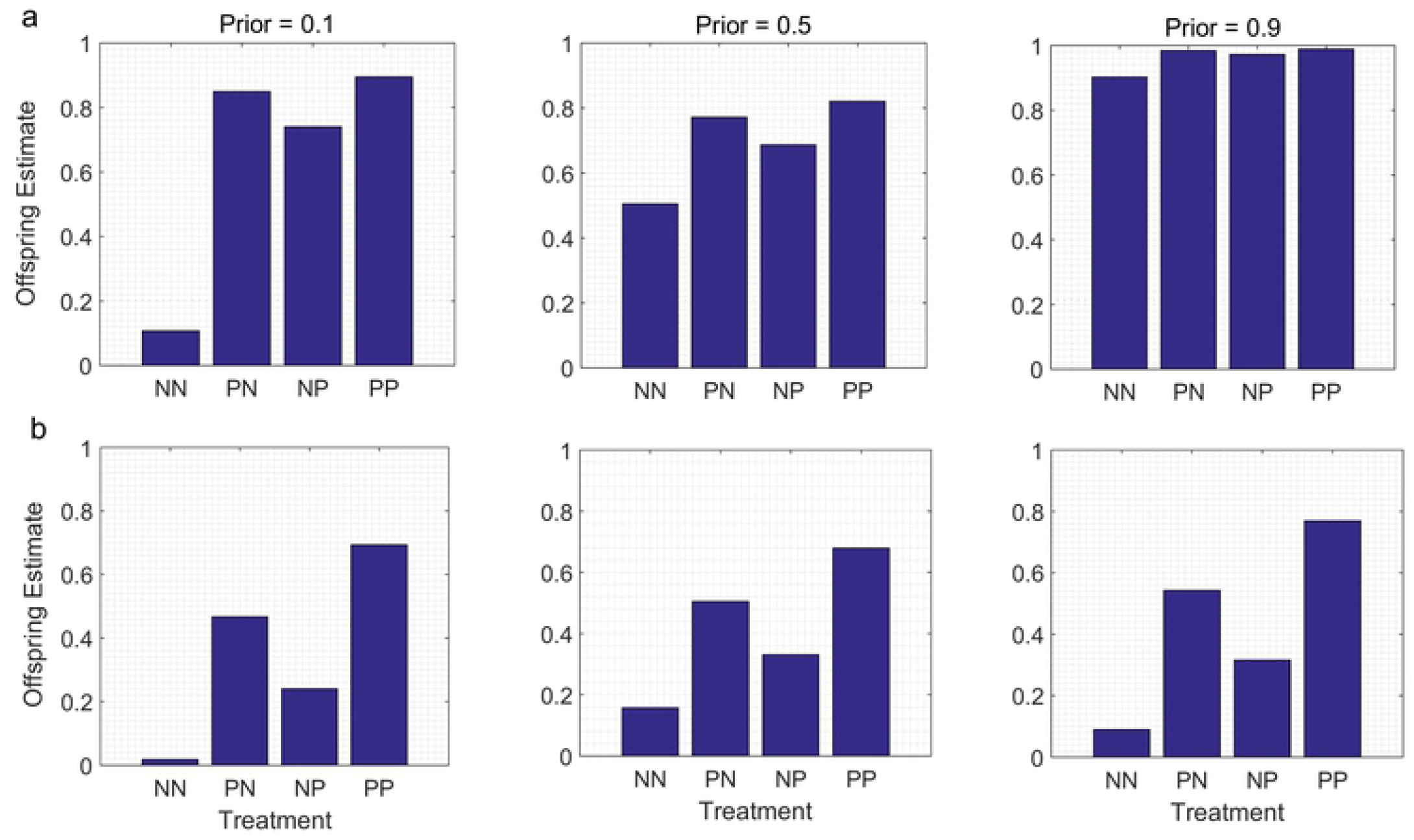
Duration of exposure to the same cues is longer for parents than for offspring. In the P treatment groups, parents are exposed to the presence of cues from predators for time four intervals, but offspring are exposed to cues from predators for only two of the four time intervals (see text). Other variables are the same as in Figure 2.

The overall patterns were otherwise similar to those found in the baseline models. That is, a jump-up pattern was detectable in the N-models when the parental Prior indicated a different value of the state than the cues in the P treatment, but not when the parental Prior indicated a value of the state similar to the value indicated by the cues in the P treatment (Figs 2a, 3a). Similarly, similar step-up patterns were detectable across a range of parental Priors in the N* models (Figs 2b, 3b). However, for comparable sets of parameter values (i.e. for the same parental Prior, and for the same likelihood function for the cues in the P treatment), the differences between the offspring estimates for the NP and PN groups were much less pronounced for the N-models than for the N* models (compare Fig 2a versus 2b, and Fig 3a versus 3b).

When information from the parents was devalued or degraded, the models predicted lower estimates of the value of the state for the PN group than for the NP group (Fig 4). The overall patterns were otherwise similar to those in the baseline models. A jump-up pattern was detectable in the N-models when the parental Prior indicated a different value of the state of the environment than did the cues in the P treatment, but not when the parental Prior indicated a value similar to that indicated by the cues in the P treatment (Fig 4a). In addition, similar step-up patterns were detectable for all parental Priors in the N* models (Fig 4b). However, for comparable sets of parameter values, the differences between the offspring estimates for PN and NP were much less pronounced for the N-models (Fig 4a) than for the N* models (Fig 4b).

**Fig 4.**
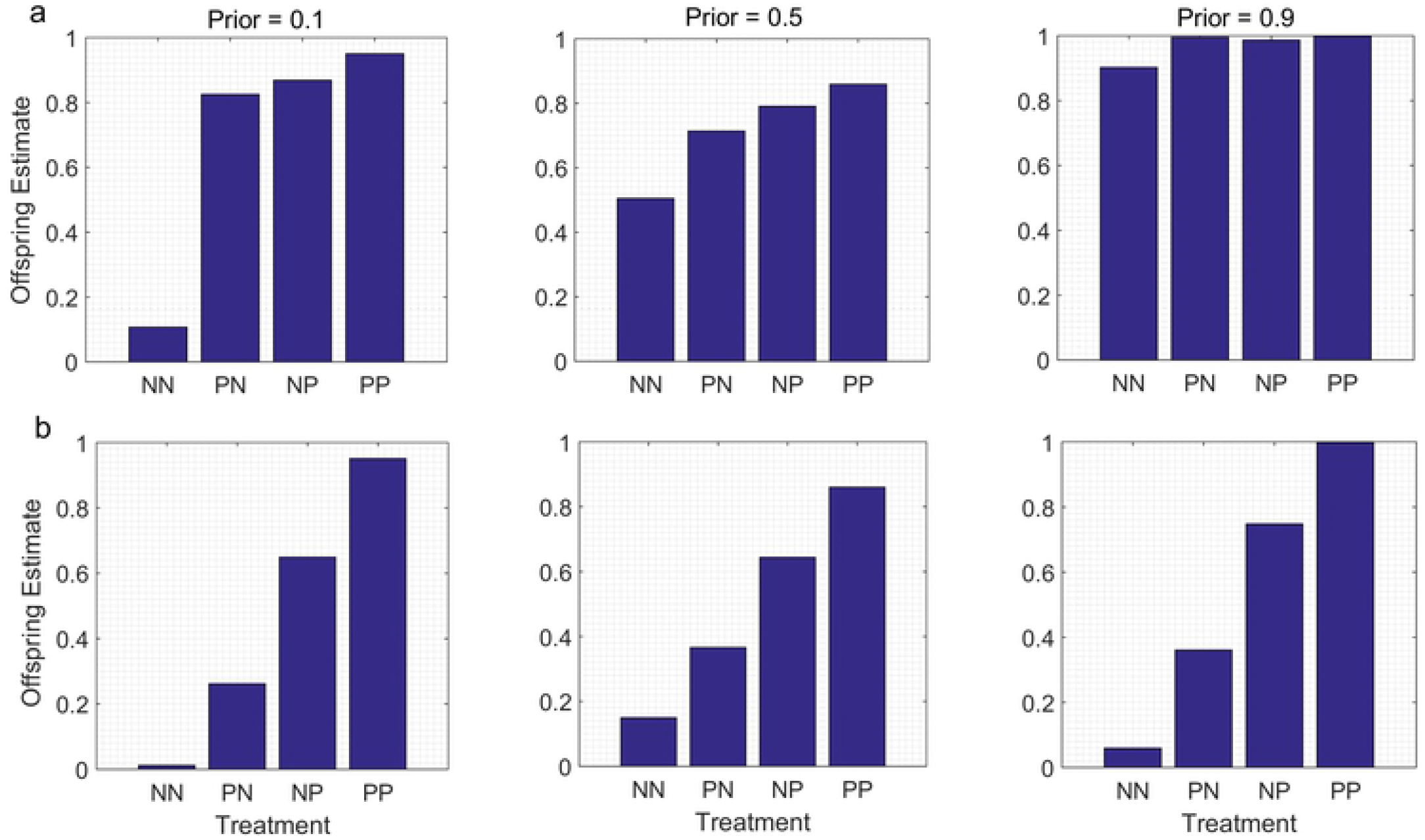
Information from parents is devalued or degraded before being passed to the offspring. We assume that parents and offspring in the P groups are exposed to the same cue for the same period of time, with a likelihood function indicated by α =8, β = 1. However, the signal provided by the parent to the offspring as a result of the parent’s exposure to the cue is less reliable; it is modelled by a likelihood indicated by α = 3.5, β = 0.44. a. N- model: information provided by conditions in the N treatment is much less reliable than the information provided by conditions in the P treatment (likelihood for conditions in the N treatment modelled by α = 1, β = 1). b. N* model: conditions in the N treatment provide information that is as reliable as conditions in the P treatment (likelihood for N modelled by α = 1, β = 8). However, the signal that parents provide to their offspring as a result of exposure to conditions in the N treatment is less reliable, with a likelihood modelled by α = 0.44, β = 3.5.

## Discussion

The current study suggests that empiricists should not be surprised to observe considerable variation among experiments in the results of studies of trans and within generational plasticity (TWP) in response to informative cues. Our results show that the offspring estimates that are assumed to provide the foundation for the offspring trait values in such experiments are expected to vary as a function of many factors, including 1) parental Prior distributions, 2) the relative reliability of the information provided in the different treatments, 3) differences in the duration of exposure to the same cues for parents and offspring, 4) the extent to which information based on cue-exposure for parents is devalued, relative to the information based on cue-exposure for offspring, and 5) the extent to which information passed from parents to offspring is degraded, relative to the information based on the personal experience of the offspring. We illustrate these findings by modelling fully factorial experiments in which parents, offspring, both or neither are exposed to cues from predators (in P treatments) or are not exposed to those cues (N treatments).

Our results confirm previous results showing that interactions between parental Priors and relative cue reliability are expected to have a major impact on the patterns of offspring information states in empirical studies of TWP [25]. We analyzed two extreme situations for differences in the reliabilities for the two treatments: N-models, in which the information provided by conditions in the N treatment is much less reliable than the information provided by conditions in the P treatment, and N* models, in which the information provided by the conditions in the P and the N treatments is equally reliable. A possible example of the former might be an experiment in which the cues from the predator in the P treatment consists of a single, near-escape from a predator over a period of several months. In this case, conditions in the P treatment might provide a reliable indication that the predator in question lives at the current locality, whereas the conditions in the N treatment may provide a less reliable indication that it may not. By way of analogy, being the victim of a robbery once over the course of a year might indicate with a high degree of reliability that thieves are active in your neighborhood, but if such incidents rarely occur, not being robbed over the course of a year need not indicate with the same level of reliability that they are not. An example of a situation in which the conditions in the P and the N treatments might provide equally reliable information is when the subjects in the P treatment are exposed for an extended period to a high concentration of kairomones from a predator and the subjects in the N treatment are maintained in the absence of kairomones for the same period of time. In this situation, investigators typically assume that the conditions in the P treatment reliably indicate that predator density is high, whereas the conditions in the N treatment indicate with a comparable level of reliability that predator density is low, since the concentration of kairomones never exceeded the preys’ threshold for detection (see [25] for additional discussion).

The current results confirm previous results indicating interactions between parental Priors and the relative reliability of the cues in the different treatments (here P vs N) are expected to have major effects on the patterns of offspring estimates in studies of TWP in response to informative cues. In particular, the predicted patterns of offspring estimates were highly dependent on parental Priors in the N-models, but virtually independent of parental Priors in the N* models [25].

These results imply that if investigators use an experimental protocol in which the information provided by the different treatments differs in its reliability (as in the N-models), the results of their experiments might vary as a function of the population-of-origin or the genotypes of their subjects. This follows from the assumption that subjects from different populations might have different parental Priors, based on information from their ancestors, e.g. from genes, inherited epigenetic factors or grand-parental effects. In addition, if the parents for an experimental study of TWP were collected from the wild, then variation among populations in parental experiences earlier in life could also contribute to variation among populations in parental Priors. Along the same lines, different genotypes from the same population might begin with different prior estimates of the value of the same state of the environment [e.g. 24], so that the results of studies of TWP based on clonal organisms might vary depending on the clone that was selected for the study.

Indeed, if investigators used an experimental protocol in which the conditions in the different treatments differed with respect to the reliability of their information (as in the N-models), the choice of population or genotype for a study might determine whether one could even detect plasticity. For instance, in all of the N-models analyzed in this article, a ‘jump-up’ pattern (NN < PN,NP < PP) for offspring estimates was observed when the value of the state of the environment indicated by the parental Prior strongly contradicted the value of the state indicated by the cues in the P treatment. However, when the value of the state indicated by the parental Prior was similar to the value indicated by the cues in the P treatment, the offspring estimates were virtually identical in all four treatment groups (NN≈ PN≈ NP≈ PP). In the latter situation, one would not expect to observe either WGP or TGP.

In contrast, consider a situation in which an investigator used an experimental protocol in which the cues in the different treatments were all equally reliable (as in the N* models). In this case, our results suggest that the choice of a population or a genotype for the experiment might not matter. As we show here for all of the models, in this situation we expect to observe a similar step-up pattern (NN < PN,NP < PP) regardless of the parental Prior.

As intuition would suggest, the main effect of assuming that cue duration differed for parents and offspring or that the information provided by parents was either devalued or degraded was to generate different values of the offspring estimate for the PN and the NP treatments; in the absence of these assumptions, the offspring estimates were the same for the PN and NP treatments. However, for the same parental Prior and for the same cues in the P treatment, the effects of differences in cue duration or the devaluation or degradation of information from parents were much more pronounced in the N* models than in the N-models. This suggests that in general, it would be easier to detect the effects of cue duration or of the parental devaluation/degradation of information on offspring estimates or trait values in TWP studies if one used an experimental protocol in which the information provided by both treatments was similarly reliable than if one used a protocol in which the information provided by one treatment was much more reliable than the information provided by the other.

If we just focus on situations in which the information provided by the conditions in both treatments is equally reliable (the N* models), the results indicate, as intuition would suggest, that the effects of those exposures on offspring estimates would be stronger for the generation that was exposed to the cues for the longer period of time. This result supports earlier suggestions that under natural conditions, parental experiences might have a stronger impact than offspring experiences on the phenotypes of offspring during the juvenile period, because the cumulative experiences of parents over their lifetimes provides more reliable information than is provided to offspring based on their own experiences since conception [35, 36]. As a practical matter, these results indicate that variation among experimental studies of TWP in the relative duration of exposure to the same cues for parents and offspring would, all else being equal, be expected to contribute to variation in their results.

The models of information-updating presented here may also provide a useful point of departure for investigating other aspects of the timing of cue-exposure that might contribute to the variation results observed in empirical studies of TWP. For instance, the models analyzed herein ignore sensitive periods, i.e. situations in which exposure to the same cues have different effects on trait values, depending on the age of onset of the cue-exposures [37-40]. Although traditionally the literature on sensitive periods has focused on the effects of an individual’s own experiences early in life on its trait values later in life (i.e. WGP), recently researchers have begun to consider the possibility of sensitive periods for TGP, i.e. situations in which exposure to cues for parents have different effects on the trait values of their offspring, depending on the age at which those exposures occurred in the parents, or the age at which the offspring received the information from their parents [e.g. 26, 30, 41]. However, at this point it is unclear how sensitive periods of parents, offspring or both would interact to affect offspring estimates of conditions in the external environment.

As a general rule, we would expect the effects of sensitive periods on offspring information-states to depend on the protocol used in a given experiment. Personal exposure to cues in the external environment can’t begin to affect the information-states of offspring until the offspring have become capable of detecting those cues (e.g. as embryos, newborns, or hatchlings). An additional complication is that in order to demonstrate that cue-exposure for parents affects the trait values of their offspring, exposure to cues in the external environment for parents must end before their offspring become capable of detecting the cues on their own. Hence, if both parents and offspring were continuously or repeatedly exposed to the cues of interest from the embryo stage until just prior to offspring production, both generations would be exposed to those cues throughout their sensitive periods, no matter if or when they occurred in either generation. In that case, the predicted patterns of offspring estimates would be the same as those predicted in the absence of sensitive periods (e.g. see Figs 1 and 4). However, if parents or offspring had sensitive periods for the effects on cue-exposures on offspring trait values, if these periods occurred at different ages in the two generations, and if exposure to the inductive cues was limited to restricted periods in both generations, then the effects of sensitive periods on offspring estimates are unclear. This would be a worthwhile topic for further empirical and theoretical study.

Our results also confirm the intuitive notion that when information from parents is devalued relative to information from offspring, exposure to cues for parents would have a weaker effect on offspring estimates than equivalent exposure to the same cues in the offspring. Theory predicts that the devaluation of information from parents should occur as an evolved, adaptive response to reduced levels of autocorrelation between parental environments and offspring environments (see Introduction). In other words, the devaluation of information from parents is expected when, under natural conditions, the value of a state of the environment is likely to change between the time that parents are exposed to cues and the time that their offspring are exposed to the same cues. For instance, if we assume that the true value of the state gradually changes over time, the devaluation of information from parents would be inversely related to the amount of time that elapsed between the time that parents were exposed to cues and the time that their offspring were exposed to the same cues. In turn, this implies that for subjects from the same population, the devaluation of information from parents would be less pronounced in an experiment in which parents were exposed to the cues of interest just before their offspring were conceived and their offspring were exposed to the same cues soon after hatching than in an experiment in which both parents and offspring were exposed to the cues soon after hatching.

Of course, in natural populations, the true value of a state of the environment need not change gradually over time, but may instead occur at specific times, lifestages or life-history landmarks. For instance, in a species in which the value of a state (e.g. predator density) does not change over the distances typically traveled by natal dispersers, information based on a parent’s exposure to kairomones from a predator in its natal habitat would be also be relevant to its offspring’s estimate of predator density in its own natal habitat. In contrast, in a species in which predator density varies over typical dispersal distances, information based on a parent’s exposure to kairomones in its natal habitat would be less useful for estimating the predator density in its offspring’s natal habitat. In other words, theory suggests that we should expect the devaluation of information from parents to differ among different populations or species, depending on the extent to which the environmental factors of interest varied spatially over the distances relevant to natal dispersal, or varied temporally within and across generations. Hence, one could potentially test predictions about the devaluation of information from parents using comparative data from field studies indicating whether, how, and when particular environment conditions of interest are spatially and temporally correlated for parents and their offspring. Unfortunately, at present this data is still sparse [42-44].

In addition, our results also indicate that if information from parents is degraded before it is passed along to their offspring, exposure to cues for parents would have a weaker effect on offspring estimates than equivalent exposure to the same cues in offspring. However, in contrast to the duration of exposure to cues, which can be readily manipulated or controlled by investigators, the degradation of information from parents cannot. The degradation of information from parents is assumed to occur as a non-adaptive consequence of the proximal mechanisms by which parents transfer information to their offspring. That is, the degradation of information from parents to their offspring is assumed to be a result of the ‘noise’ introduced into the flow of information from one generation to the next, based on the series of processes that intervene between the time that mothers and/or fathers are personally exposed to the cues and the time that an offspring receives signals from their parents based on the parent’s estimate of the value of the state [30].

Because we assume that the degradation of information occurs as a result of unavoidable constraints in the mechanisms responsible for the transfer of information across generations, we would expect the degradation of information from parents to be phylogenetically conservative, i.e. similar for populations and species with comparable patterns of offspring development and parental care. Thus, in contrast to the situation for the devaluation of information from parents, we would not expect the degradation of information from parents to vary across populations or closely-related species, as a function of the extent to which the environmental conditions experienced by parents and offspring were correlated with one another across space and time. Instead, testing hypotheses about the effects of the degradation of information from parents to offspring will require detailed information about the proximal mechanisms which mediate the flow of information from parents to offspring in a given taxon. At present, little is known about this topic; this would be another fruitful topic for future research.

Under certain conditions, models which predict patterns of offspring estimates may help shed light on the patterns of offspring trait values observed in empirical studies of TWP. One important condition is that the inductive cues only provide information, as opposed to also having direct, long-lasting effects on the somatic states of parents, offspring or both. Detailed discussion of the difference between inductive experiences which affect development because they provide information and inductive experiences which affect development because they have persistent effects on somatic states can be found in [6, 9, 45, 46]. Examples of information-only inductive experiences include stimuli produced by predators, conspecifics, or competitors. Examples of inductive experiences which might provide information, but which also have direct effects on a developing organism’s somatic state, include food deprivation in animals, shade in plants, and extreme temperatures in either. As Nettle and Bateson (2015) point out, if an inductive experience is information-only, one can imagine a single loss-of-function mutation that abolishes an individual’s ability to detect that cue, but which leaves the developing individual otherwise unaffected. In contrast, if an inductive experience has a direct and lasting impact on an individual’s state, one would expect that experience to affect its trait values even if the individual was unable to sense that it had that experience. Although it is possible to make general predictions about how particular trait values might change in response to exposure to information-only cues over the course of development [e.g. 17, 20], this is much more difficult when experiences also have direct, lasting effects on an individual’s metabolism, growth or other aspects of its somatic state (but see 16).

A second important condition for assuming that offspring estimates will be directly related to the expression of a given trait in the offspring is evidence that the trait is advantageous when the offspring is in the environment indicated by the cues in the experiment. For example, in empirical studies of TWP in response to cues from predators, investigators often focus on inducible defenses, e.g. behavioral or morphological traits which have been shown to increase survivorship when offspring are in the presence of predators. In such cases, it is reasonable to assume that those traits would be more strongly expressed if a subject’s estimate of the value of an environmental state that contributes to predation risk (e.g. predator density) was high than if it was low. In contrast, analyses of offspring information-states are much less useful for predicting the plasticity of traits for which the expected adaptive response to different information states is unknown or uncertain.

However, if a cue is information-only, if the adaptive phenotypic response to that cue is clear, and if one has a reasonable idea about the relative reliability of the information provided by the conditions in the different treatments, then these models can provide a useful benchmark against which to compare the patterns for offspring trait values observed in empirical studies of TWP. For instance, if parents and offspring are both exposed to the same cues from predators from birth to maturity, if one measures inducible traits known to improve survivorship in the presence of that predator, and if it is reasonable to assume that the cues in the P treatment and the cues in the N treatment are equally reliable, then the models in the current paper predict a ‘step-up’ pattern for offspring trait values, in which the values for an antipredator trait for the PN group are either the same or lower than the values for the NP group (see Figs 1b and 4b). In addition, if information from the field indicates that the environmental state of interest might change between the parental and offspring generations, one would expect the trait values for the NP group to be higher than the trait values for the PN group.

An example of an experiment for which these conditions appear to be satisfied is a classic study of TWP in which *Daphnia cucullata* parents and offspring were exposed to kairomones from a predator (*Chaoborus flavicans*) from birth to first reproduction, and then relative helmet length was measured at the age of first reproduction in the offspring [2]. Other experiments have shown that kairomones from *C. flavicans* induce the development of larger helmets in *D. cucullate* [47], and that relatively large helmets protect juvenile and adult *D. cucullata* from this predator [48]. In addition, field studies of *C. flavicans* indicate that the risk it poses to *Daphnia* spp. varies within seasons across the temporal scales that might encourage the devaluation of information from parents [49]. Finally, continuous exposure of kairomones from a predator from birth to maturity in a P treatment is typically assumed to provide reasonably reliable information about the density of that predator, while the absence of cues from the predator from birth to maturity in an N treatment is assumed to provide comparably reliable information indicating that the density of that predator is low. As predicted by the models described herein under this set of conditions Agrawal et al. [2] reported a step-up pattern (NN < PN < NP < PP), in which the trait values for all four groups were significantly different from one another.

Finally, the current study shows how an appreciation of the ways that information from different sources is expected to combine within and across generations reveals a number of questions that investigators might want to consider when they are planning or interpreting the results of empirical studies of TWP in response to inductive cues. These are summarized as follows:

1. Are the inductive cues information-only, or could the inductive experiences also have direct, lasting effects on the somatic states of either the parent or their offspring? Insights from models of development based on information-updating are currently most useful for predicting trait values when cues are information-only.
2. Are the conditions in the different treatments likely to provide equally reliable information about a state of the environment, or are the conditions in one treatment likely to provide much more reliable information than those in the other treatment? As was shown here and in [25], we expect the patterns of offspring estimates in TWP studies to dramatically differ in these two situations.
3. Are parents and offspring exposed to the same cues for the same period of time? As is shown in the current article, all else being equal, differences in the duration of cue-exposure for parents and offspring are expected to affect the patterns of offspring estimates, especially if the cues in the different treatments are equally reliable.
4. Is there strong existing support for the assumption that a particular response in a particular trait to a particular cue is likely to be adaptive? In such cases, it is more likely that differences among the treatment groups in offspring information states at the end of the experiment will be related to differences among those groups in the trait values expressed by the offspring at the end of the experiment.

In addition, our results indicate that it may not be necessary to invoke assumptions about developmental constraints, costs of sampling, or the fitness consequences of trait values of offspring to account for at least some of variation in results observed in empirical studies of TWP. Instead, we have shown that even if we restrict ourselves to experiments with information-only cues, we should expect to observe considerable variation in their results, as a function of variation in parental priors, the reliability of the cues in the different treatments, differences in the duration of cue-exposure for parents and offspring, and the extent to which information from parents was either devalued or degraded in comparison to information from offspring.

## Acknowledgments

We thank Yifeng Xu for help with programing. This material is partially based upon work supported by the National Science Foundation under Grant No. IOS 1121980 and the National Institutes of Health under award number 2R01GM082937-06A1.

